# The rate of identical-by-descent segment sharing between close and distant relatives

**DOI:** 10.1101/2025.07.30.667761

**Authors:** Amy L. Williams

## Abstract

Genetic relatives share long stretches of DNA they co-inherited from a common ancestor in identical-by-descent (IBD) segments. Because children inherit half their parents’ genomes, the expected amount of DNA relatives share drops by 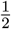 for each generation that separates them, being 2^−*d*^ for *d*-degree relatives. Even so, there is substantial variance in sharing rates, such that most distant relatives share zero IBD segments. We characterized IBD segment sharing between relatives by simulating 100,000 pairs for each of first through eighth cousins, including once-removed and half-cousins, while modeling both crossover interference and sex-specific genetic maps. Our results show that 98.5% of third cousins share at least one IBD segment, while only 32.7% of fifth cousins and 0.961% of eighth cousins have such sharing. These sharing rates are substantially higher than those that arise from models that ignore the more elaborate crossover features. The resulting segment count distributions are available with an interactive segment length threshold at https://hapi-dna.org/ibd-sharing-rates/.

## Introduction

Relatives share one or more common ancestors, a heritage that manifests in some relatives as long shared segments of DNA or identical-by-descent (IBD) segments ^1^. Sharing even a distant ancestor with another individual carries the possibility of the two sharing one or more IBD segments, but the average amount of DNA relatives share decreases with the number of generations that separate them. A parent-child pair shares 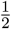 of their genomes IBD, and this factor of two—which halves the DNA transmitted across every generation on the lineage that connects two individuals—implies that on average relatives separated by *G* generations share a fraction 2^−*G*^ of their genomes in IBD segments (for one common ancestor) or 2^−*G*+1^ (for descent from an ancestral couple). Throughout, we use IBD to mean either identical-by-descent or identity-by-descent depending on context.

Meiosis constrains parent-child pairs to share exactly 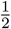 of their genomes IBD, with no variance, but due to the random nature of recombination and finite genomes, the amount of DNA shared for every other relationship does vary. More specifically, crossovers occur in each generation, but their rate is roughly 1.2 × 10^−8^ per base-pair per generation in humans ^2^, so non-recombined segments transmitted from one generation to the next are large, averaging roughly 83 Mb (assuming a Poisson crossover distribution, which corresponds to an exponentially distributed distance between crossovers). This, coupled with independent segregation, or a 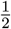 probability of any non-recombined segment being transmitted, implies that distant relatives can easily share zero IBD segments.

Characterizing the rate at which relatives share IBD is important for a range of analyses, as IBD sharing is central to IBD-based association testing ^3^ and to approaches for estimating heritability without environmental bias ^4^. Moreover, IBD sharing rates are fundamental to detecting and inferring relatedness among individuals ^5^, and direct-to-consumer (DTC) genetic testing companies use IBD segments for this purpose ^6^. The popularity of these tests has led millions of individuals to learn of their close and very distant relatives, which has helped fuel family history research (i.e., genetic genealogy).

Studying the distribution of IBD sharing between relatives is stymied by a lack of data for known relatives. Few genetic studies collect information about relationships between the participants, and though some datasets with many moderately sized pedigrees exist ^7,8^, data for large pedigrees (which include pairs of distant relatives) are less abundant ^9^. Even if data for many pairs of purported relatives were collected, such as might be feasible for DTC companies, challenges of incorrectly or incompletely reported relationships would cloud any analysis. Furthermore, pedigree collapse ^10,11^—whereby individuals are descended from ever increasing fractions of the population for each earlier generation going back through time—implies that we are in fact related in myriad ways, making the true ancestor that transmitted an IBD segment difficult to discern. As well, IBD segment detection is error prone—with both false positives and false negatives— especially for short segments.

An alternative to empirical data analysis is using theoretical models to determine the rate at which various kinds of relatives share IBD segments. One classic approach by Donnelly used continuous-time Markov chains and a group theory-based collapsing of states to calculate these probabilities ^12^. As described in Results, the rates from that work closely match our simulated data—but only when ignoring sex-specific genetic maps and crossover interference. These latter processes pose significant challenges for incorporating into analytical or numerical methods ^13^; yet simulating with these processes yields better fits to IBD sharing from real data ^13^, and their presence has a sizeable impact on IBD sharing rates (Results). Furthermore, minimum segment length thresholds are key factors in analyses of real data, but it is not clear how to impose such thresholds using Markov chains.

To characterize the distribution of IBD sharing among relatives unencumbered by the important but complex factors inherent in real data analysis, while also accounting for more elaborate features of recombination, we simulated first through eighth cousins, including once removed relatives and half-cousins (30 relationships total). Because the IBD segments are generated by the simulator, they necessarily descend only from the common ancestor(s) that define the relationship and they are not subject to detection error. The result is a set of highly precise IBD segment distributions for every relationship type, with even 1 bp long segments recorded, and also distributions that only include segments longer than several thresholds.

## Methods

We used Ped-sim ^13^ to simulate 100,000 relative pairs for each relationship and included both crossover interference modeling ^14,15^ and sex-specific genetic maps ^2^ to obtain a realistic representation of meiosis ^13^. The sexes of all individuals in the simulated pedigrees are male or female with equal probability, thus capturing the heterogeneous spectrum of descent underlying real relatives. Separately, we simulated another set of 100,000 pairs for each relationship using a Poisson crossover distribution and a sex-averaged genetic map, mirroring Donnelly’s model ^12^.

The simulated relationships include: full first through eighth cousins, abbreviated below as *N* C for *N* th cousins (e.g., 1C for first cousins); full first cousins once removed through seventh cousins once removed, abbreviated as *N* C1R for *N* th cousins once removed; half-first through half-seventh cousins, abbreviated h*N* C for half-*N* th cousins; half-avuncluar (e.g., a half-uncle-niece pair), abbreviated hAV; and half-first cousins once removed through half-seventh cousins once removed, abbreviated h*N* C1R for half-*N* th cousins once removed.

## Results

The relationships we consider span 3rd through 17th degree, with one full cousin type and one half-relationship for each degree. For example, full first cousins and half-avuncular pairs are both 3rd degree relatives, while first cousins once removed and half-first cousins are each 4th degree relatives. Relatives of the same degree share the same average amount of DNA, and the rates that full and half-relatives of the same degree share at least one IBD segment with each other are also similar, with a few notable differences described below.

Figure 1A depicts the percent of relatives that share any IBD, incorporating segments of any length, with colors indicating the number of segments shared (the plots here give rates for full relatives; plots for half-relatives are available on the website). All 3rd through 5th degree relatives (1C, 1C1R, 2C, hAV, h1C, and h1C1R pairs) share at least one IBD segment (and in nearly all cases ≥ 5 segments), but 6th degree relatives have a small chance of sharing no IBD—0.07% for 2C1R and 0.16% for h2C. The probability of sharing one or more segments is *>*97% for 7th degree (3C and h2C1R) and closer relatives, but drops rapidly as the relationships grow more distant. For instance, 10th degree relatives have IBD sharing between ∼50% of pairs (51.3% of 4C1R and 48.4% of h4C), and even fewer relatives share IBD for each more distant degree. The sharing probability drops by less than 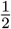 for each additional degree because even some distant relatives share ≥ 2 IBD segments, thus increasing the probability that a descendant—corresponding to a one-degree more distant relationship—will inherit at least one segment. Additionally, even when a parent shares only one segment with a relative, if the parent transmits a crossover within that segment, the child will necessarily inherit part of it.

**Figure 1.**
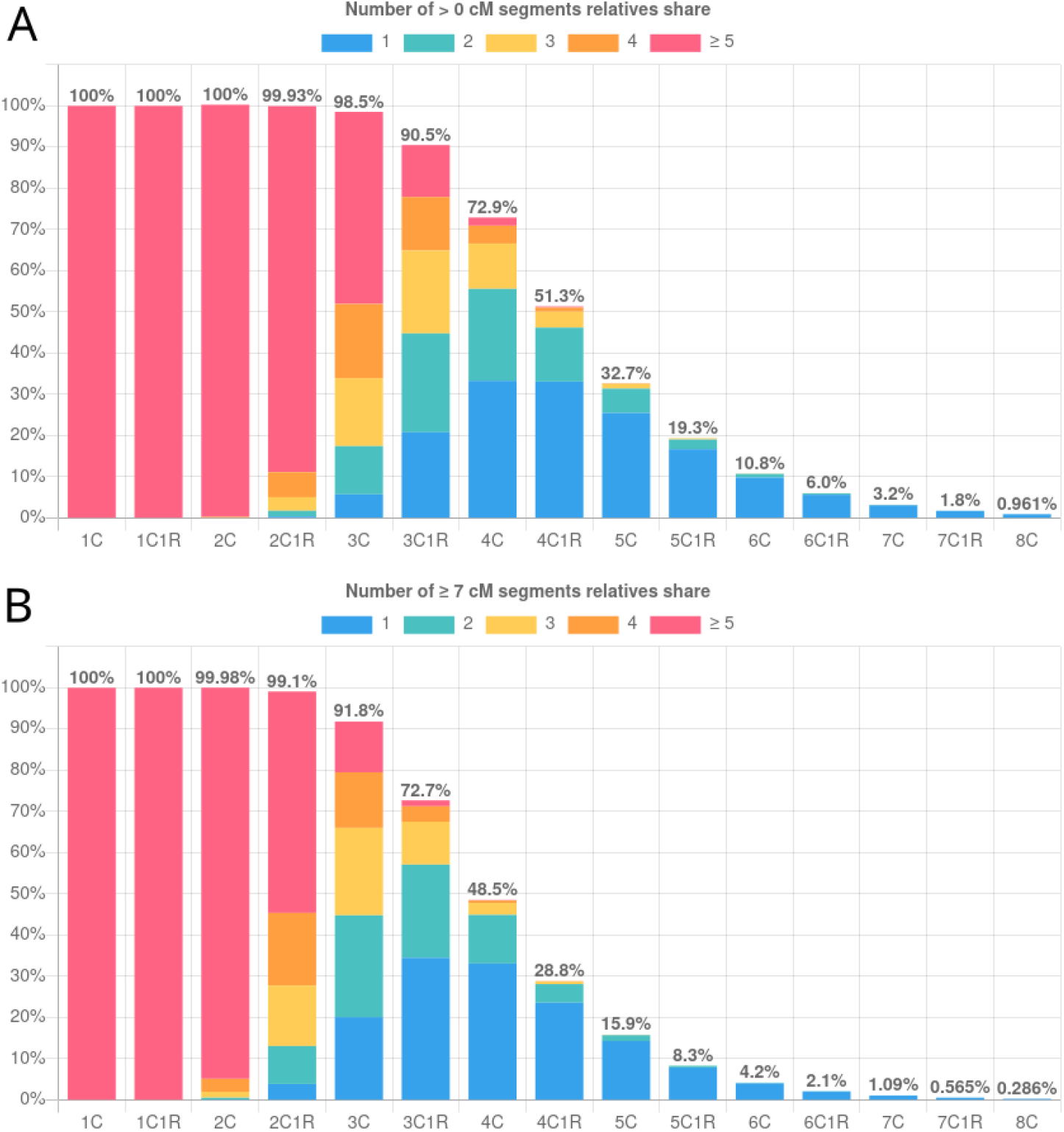
Percent of relatives that share one or more IBD segments. Plots show sharing rates between relative pairs for all full relationships we simulated, considering (A) segments of any length and (B) segments ≥ 7 cM. Abbreviations are *N* C for *N* th cousins and *N* C1R for *N* th cousins once removed. Plots for half-relatives and other segment length thresholds are available at https://hapi-dna.org/ibd-sharing-rates/. The web-based plots also provide the percent of relatives that share the indicated numbers of segments via a tooltip pop up.

Compared to Donnelly’s exact calculations, if (as in that work ^12^) we leverage a sex-averaged genetic map and Poisson crossover placement (i.e., ignore interference), the probabilities derived from both approaches are consistent, with a mean absolute difference per relationship of 0.24% (maximum 0.66%) (Supplementary Table 1). However, simulating with sex-specific maps and interference yields meaningfully higher IBD probabilities, with the 10th degree 4C1R sharing IBD in 51.3% of pairs when including these features and only between 48.6% of pairs without them (2.7% absolute difference). The magnitude of the difference between the models varies by degree of relatedness: there is no difference for 3rd and 4th degree relatives (where all pairs share IBD under both models); a maximum of 3.1% more relatives carry IBD for the 9th degree h3C1R; and only 0.076% more carry IBD between the 17th degree h7C1R (Supplementary Table 1). While the difference for distant relatives is small in absolute terms, it is large in relative terms with, e.g., 12.3% more 8C sharing IBD under the more realistic model. The difference between the models is primarily driven by crossover interference, as relatives simulated using sex-specific maps and Poisson localization have very similar IBD sharing rates to simulations that use a sex-averaged map (using random ancestor sex assignment; not shown).

Overall, full relatives have non-zero IBD sharing somewhat more often than half-relatives of the same degree—e.g., 32.7% of 5C share at least one IBD segment compared to 30.4% of h4C1R. Full relatives have undergone one more round of meiosis than half-relatives of the same degree but share two common ancestors instead of one; e.g., there are four meioses on the two lineages connecting first cousins but only three meioses separate half-avuncular pairs. The consequences of more meioses with two common ancestors are that full relatives generally share a larger number of segments that are smaller than those of the corresponding half-relatives (and thus there is a higher probability of full relatives inheriting at least one segment).

We made these IBD sharing distributions available online at https://hapi-dna.org/ibd-sharing-rates/, which has an option to select a minimum segment length. Because IBD segment detectors have minimum length thresholds, very short segments are unlikely to be detected, motivating the analysis of only segments longer than a specified length. When considering segments ≥ 3 centiMorgans (cM), the distributions shift, such that, e.g., only ∼40% of 10th degree relatives share any IBD (40.9% of 4C1R and 39.2% of h4C). With a ≥ 7 cM threshold, these same relatives share IBD between only ∼28% of pairs (28.8% of 4C1R and 28.2% of h4C), while 9th degree relatives have IBD sharing between ∼48% of pairs (48.5% of 4C and 47.9% h3C1R) (Figure 1B).

The most extreme segment length threshold we analyzed is ≥ 40 cM, and at that threshold, even a few 3rd degree relatives (0.29% of 1C and 0.08% of hAV) do not inherit any sufficiently long segments. Because half-relatives generally share fewer but longer segments, in fact, more half-relatives have at least one ≥ 40 cM segment compared to full relatives of the same degree. For example, between 5th degree relatives, 55.1% of 2C and 65.0% of h1C1R share at least one ≥ 40 cM segment.

The average amount of DNA 10th degree relatives are expected to share is 2^−10^ of their genomes, or about 6.5 cM in the genetic maps we used (sex averaged autosomal length 3,346 cM; total diploid length 6,693 cM) ^2^. This expectation includes relatives that share zero IBD and matches our observations (mean 6.5 cM shared considering all 4C1R). Yet conditional on sharing at least one segment, on average 4C1R share 12.7 cM of DNA, meaning that these 10th degree relatives have sharing closer to the expectation for 9th degree relatives. An even more extreme case is the 15th degree 7C who, conditional on carrying IBD, share an average of 6.34 cM, which is roughly in line with 10th degree relative sharing. This shift in IBD amounts for relatives that share at least one IBD segment can lead to substantial bias in relatedness estimates, as Jewett recently pointed out and developed models to correct for ^16^.

## Discussion

The IBD segment distributions we simulated are informative for a host of analyses, including genetic genealogy research to aid in determining the relationship between individuals—or the probability that two presumed relatives may not share any IBD. Although we included only 30 relationships, in fact, these IBD rates are equivalent to a vast number of other relationship types: those who are separated by the same number of generations and have the same number of shared ancestors. For example, first cousins twice removed (a given person and their grandparent’s first cousin) share two common ancestors and are separated by six generations, which is the same as second cousins. The lineage(s) the meioses occurred on does not affect the IBD sharing rates between the relatives, so they are broadly applicable to many relationships.

An important limitation of these distributions is their simulated nature. Because empirical data will necessarily be noisy due to data quality and IBD detection limits, IBD distributions from real relatives may differ from these idealized simulations. Furthermore, although we modeled non-canonical features of meiosis, we did not include other known features, such as variable crossover rates due to genetic effects ^17^, age effects ^15^, and the covariance of crossover rates across chromosomes (the so-called gamete effect ^17^). The segments also do not reflect additional IBD that may arise due to endogamy, in founder populations, or more generally from pedigree collapse. Even so, as a general guide to IBD sharing distributions among relatives, these rates are finding widespread use among genetic genealogists and are also a useful resource for population geneticists ^16^.

## Acknowledgements

We thank Shai Carmi and Ethan Jewett for helpful feedback on this paper. This work was supported in part by National Institutes of Health grant R35 GM133805.

## Supplementary material

**Supplementary Table 1:** Percent of relatives that share one or more IBD segments under several models. The degree column lists the degree of relatedness. Percentages for both full and half-cousins are given for: Donnelly’s theoretical derivation (from Table 1 in that paper ^12^) (Donnelly); simulated relatives using a sex-averaged genetic map and Poisson crossover placement (SA+Poiss); and simulated data using sex-specific maps and a crossover interference model (SS+intf). The Δ column gives the absolute percent more relatives with IBD sharing under the SS+intf model compared to SA+Poiss. Note that Donnelly provided four significant figures whereas we include five for our simulated data.

| Degree | Full cousins |  |  |  | Half-cousins |  |  |  |
| --- | --- | --- | --- | --- | --- | --- | --- | --- |
| | Donnelly | SA+Poiss | SS+intf | $\Delta$ | Donnelly | SA+Poiss | SS+intf | $\Delta$ |
| 3rd | N/A | 100% | 100% | 0% | N/A | 100% | 100% | 0% |
| 4th | N/A | 100% | 100% | 0% | N/A | 100% | 100% | 0% |
| 5th | 100% | 100% | 100% | 0% | 100% | 99.997% | 100% | 0.003% |
| 6th | 99.88% | 99.879% | 99.928% | 0.049% | 99.71% | 99.746% | 99.840% | 0.094% |
| 7th | 97.68% | 97.804% | 98.518% | 0.714% | 96.51% | 96.587% | 97.690% | 1.103% |
| 8th | 87.94% | 87.990% | 90.518% | 2.528% | 85.16% | 85.444% | 88.087% | 2.643% |
| 9th | 69.31% | 69.953% | 72.865% | 2.912% | 65.90% | 66.564% | 69.647% | 3.083% |
| 10th | 48.04% | 48.611% | 51.308% | 2.697% | 45.17% | 45.595% | 48.449% | 2.854% |
| 11th | 30.24% | 30.647% | 32.685% | 2.038% | 28.30% | 28.686% | 30.450% | 1.764% |
| 12th | 17.84% | 18.337% | 19.329% | 0.992% | 16.90% | 16.710% | 17.781% | 1.071% |
| 13th | 10.11% | 10.362% | 10.795% | 0.433% | 9.47% | 9.507% | 10.118% | 0.611% |
| 14th | 5.58% | 5.606% | 6.021% | 0.415% | 5.24% | 5.206% | 5.554% | 0.348% |
| 15th | N/A | 3.072% | 3.238% | 0.166% | 2.89% | 2.887% | 3.112% | 0.225% |
| 16th | N/A | 1.612% | 1.771% | 0.159% | 1.56% | 1.641% | 1.719% | 0.078% |
| 17th | N/A | 0.856% | 0.961% | 0.105% | N/A | 0.849% | 0.925% | 0.076% |

